# High-resolution maps of hunter-gatherer social networks reveal human adaptation for cultural exchange

**DOI:** 10.1101/040154

**Authors:** Andrea Bamberg Migliano, Abigail Page, Jesus Gómez-Gardeñes, Sylvain Viguier, Mark Dyble, James Thompson, Nikhill Chaudhary, Gul Deniz Salali, Daniel Smith, Janis Strods, Vito Latora, Ruth Mace, Lucio Vinicius

**Affiliations:** Department of Anthropology, University College London, London WC1H 0BW, United Kingdom.; Department of Condensed Matter Physics and Institute for Biocomputation and, Physics of Complex Systems, University of Zaragoza, 50009 Zaragoza, Spain.; School of Mathematical Sciences, Queen Mary University of London, London E1 4NS, United Kingdom.

## Abstract

Are interactions with unrelated and even unknown individuals a by-product of modern life in megacities? Here we argue instead that social ties among non-kin are a crucial human adaptation. By deploying a new portable wireless sensing technology (motes), we mapped social networks in Agta and BaYaka hunter-gatherers in unprecedented detail. We show that strong friendships with non-kin optimize the global efficiency of their social networks thereby facilitating cultural exchange, and that the adaptation for forming friendship ties appears early in development. The ability to extend networks and form strong non-kin ties may explain some human distinctive characteristics such as hypersociality and cumulative culture, and the tendency to exchange ideas with unrelated and unknown individuals in megacities and online social networks.

**One Sentence Summary:** Social networks of two hunter-gatherer groups in Congo and the Philippines reveal that friendships are an ancestral adaptation for the exchange of information and culture.

## Main Text

Humans regularly interact with unrelated and even unknown individuals in large urban centres and megacities ^1^. This pattern may be a by-product of socioeconomic factors such as economic opportunities or group augmentation ^2^. An alternative option is that living and interacting with unrelated individuals evolved as a human ancestral adaptation for the general exchange of knowledge, resources, cooperative actions ^3, 4^ and cumulative culture ^5, 6^. If strong friendships with unrelated individuals are an ancestral adaptation, they should play a prominent role in structuring social networks in extant hunter-gatherer populations, which represent the best models to human social organisation before the advent of agriculture. We deployed a new portable wireless sensing technology (motes) to record all dyadic interactions within a radius of approximately 3 meters at 2-minute intervals for 15 hours a day (05:00-20:00) over a week, in six Agta camps in the Philippines (200 individuals, 7210 recoded dyadic interactions) and three BaYaka camps in Congo (132 individuals, 3397 dyadic interactions; see Table S1 for descriptive statistics for all camp networks). This allowed us to build high-resolution proximity networks mapping the totality of close-range interactions within each camp. In hunter-gatherers (who lack technology-aided communication), close proximity is an indicator of joint activities such as foraging ^7^ parental care ^8^ and information exchange^9^ among others. Proximity maps therefore provide a detailed summary of all direct social interactions in the two forager populations at a level of detail never previously recorded.

The diversity and complexity of human cumulative culture suggests that the diffusion of information in human groups occurs through optimised social networks^10^ rather than randomly. Contemporary societies provide many examples of optimised or ‘small-world’ networks ^11^, such as online communities^12^ and the World Wide Web ^10, 13^ shown to maximise the overall efficiency of information and resource flows. In the following, we argue that optimised network structures were necessary for the evolution of cumulative culture and hypercooperation in humans and hence are also observed in small-scale hunter-gatherer populations. We show that hunter-gatherer social networks are optimised by extensive interactions with unrelated individuals, that such interactions are not randomly distributed but concentrated among a small number of ‘best friends’ that help connect family units, and finally that the tendency to form friendships beyond kin is manifested early in ontogeny, suggesting a developmental adaptation for hypersociality and cultural exchange.

### Interactions with unrelated people optimise network efficiency

Our analyses show that interactions with unrelated people optimise network efficiency. We used motes data from Agta and BaYaka camps to build weighted social networks reflecting the frequency of close-range interactions between individuals (number of times individuals were recorded at close-proximity every 2 minutes) (Fig.1A and Supplementary Fig. 1). We divided the social networks into decreasing levels of relatedness, starting from individual’s close kin (parents, children, siblings, partners), extended family (grandparents, grandchildren, aunt, uncle, niece, nephew, first cousins, parents-in-law, siblings-in-law) and non-kin (all other individuals; see Methods for details of kin categorisation). We have then calculated the contribution of each level of relatedness to network global efficiency, a measure of how well the structure of the network can facilitate information flow. Results indicate that global efficiency is optimised by the links between unrelated individuals. The pattern of interactions among close kin and extended family do not affect the global efficiency of real hunter-gatherer networks relative to comparable randomised networks (see Methods for randomisations procedure). In contrast, randomisation of non-kin relationships drastically reduces global network efficiency (Fig. 1B, and Supplementary Fig. 2 for other camps). This is true for all camps both in Congo and the Philippines (Fig. 1C). Our results suggest that the efficiency of hunter-gatherer networks relies on non-kin interactions, which establish an infrastructure for information exchanges amongst unrelated families.

**Figure 1.**
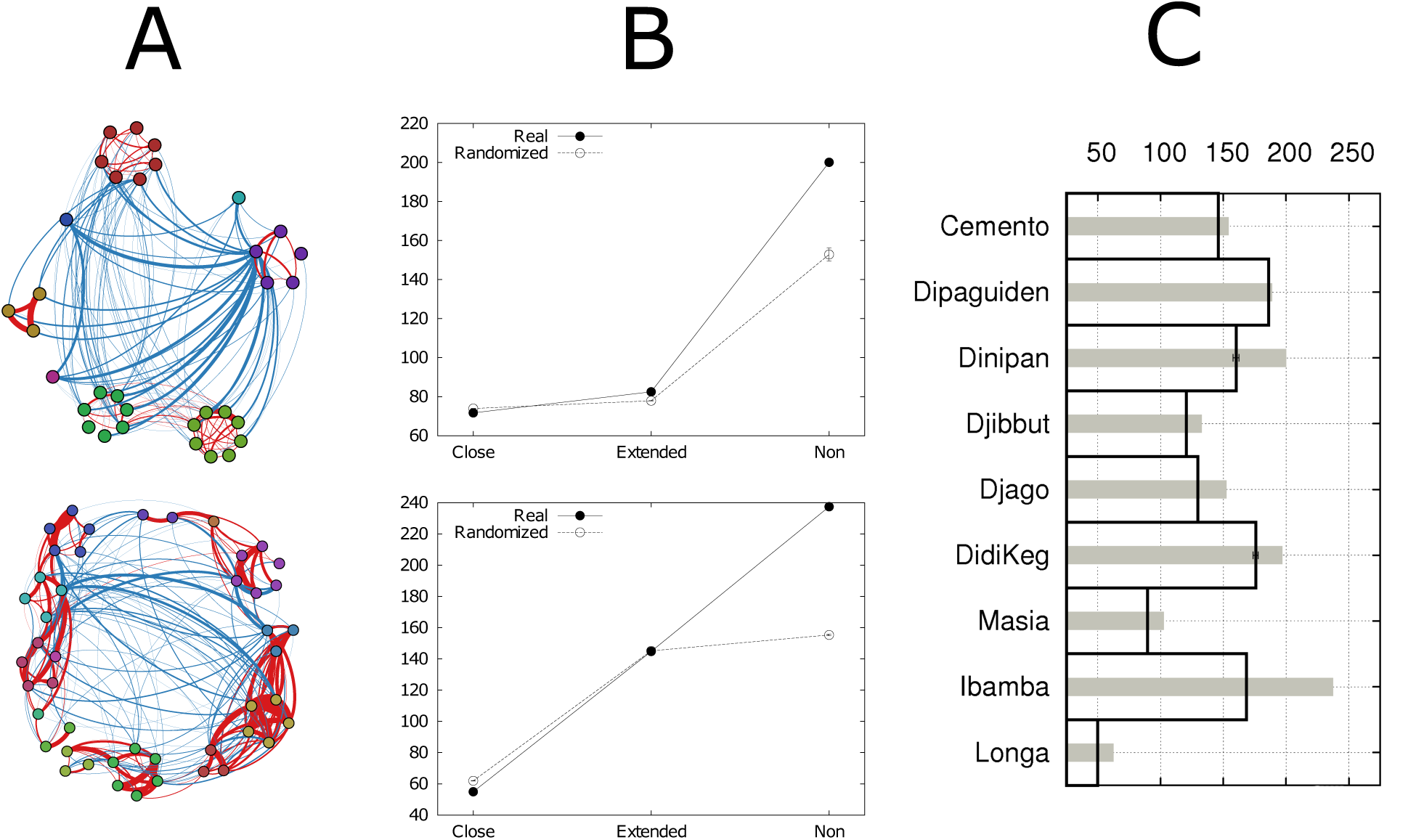
Global network efficiency depends on non-kin ties. (A) Diagrams of networks for two camps in the Philippines (top: Dinipan, N=33 people) and Congo (bottom: Ibamba, N=47 people). Nodes: individuals. Node colours: households. Red ties represent close kin or extended family, and blue ties connect unrelated individuals. Tie thickness: intensity of relationship (number of recorded close-range interactions). Graphs display the 60% strongest links. (B) Global network efficiency was compared in real (solid circles) and randomised networks of the same size and properties (open circles; see Materials and Methods for randomisation procedure). Randomisation of non-kin ties in real networks causes dramatic reduction in global efficiency, in contrast to randomisation of close kin and extended family ties. We considered averages over 1000 different randomisations. Error bars for randomisations represent standard error of mean, but are small and imperceptible. (C) Global efficiency of networks from all camps in the Philippines (top six) and Congo (bottom three). Grey bars: global efficiencies calculated from real networks. Black boxes: global efficiencies of networks with randomised non-kin ties. Error bars in black boxes: standard errors of means. In all populations, global efficiency is significantly higher in real networks than in networks with randomised ties between non-kin.

### Efficient networks require a few ‘best friends’

The frequency distribution of links among unrelated individuals in camps shows a skewed pattern. As a rule, individuals exhibit a pattern of a few intense interactions (close friends) and weaker connections with a large number of non-kin. Across all camps and age groups in both the Agta and BaYaka, people have one to four ‘best friends’ with whom they interact as strongly as with close kin (Fig. 2). In addition, measures of inequality in distribution of link weights are consistently higher among non-kin than among either close kin or extended family members. For instance, we found Gini coefficients of 0.69, 0.72 and 0.85 for close kin, extended family and non kin respectively for Dinipan, and 0.35, 0.63 and 0.92 for Ibamba (see Supplementary Table 1 for Gini coefficients in all camps). Randomisation of non-kin links, which have the effect of homogenising the strength of connections between unrelated individuals, eliminates strong friendships from networks, and significantly reduces global network efficiency. This indicates that social network optimisation does not result from a large number of random or homogeneously distributed links with all possible unrelated camp mates, but from investing in a few strong ‘best friends’ in addition to an extended net of social acquaintances.

**Figure 2.**
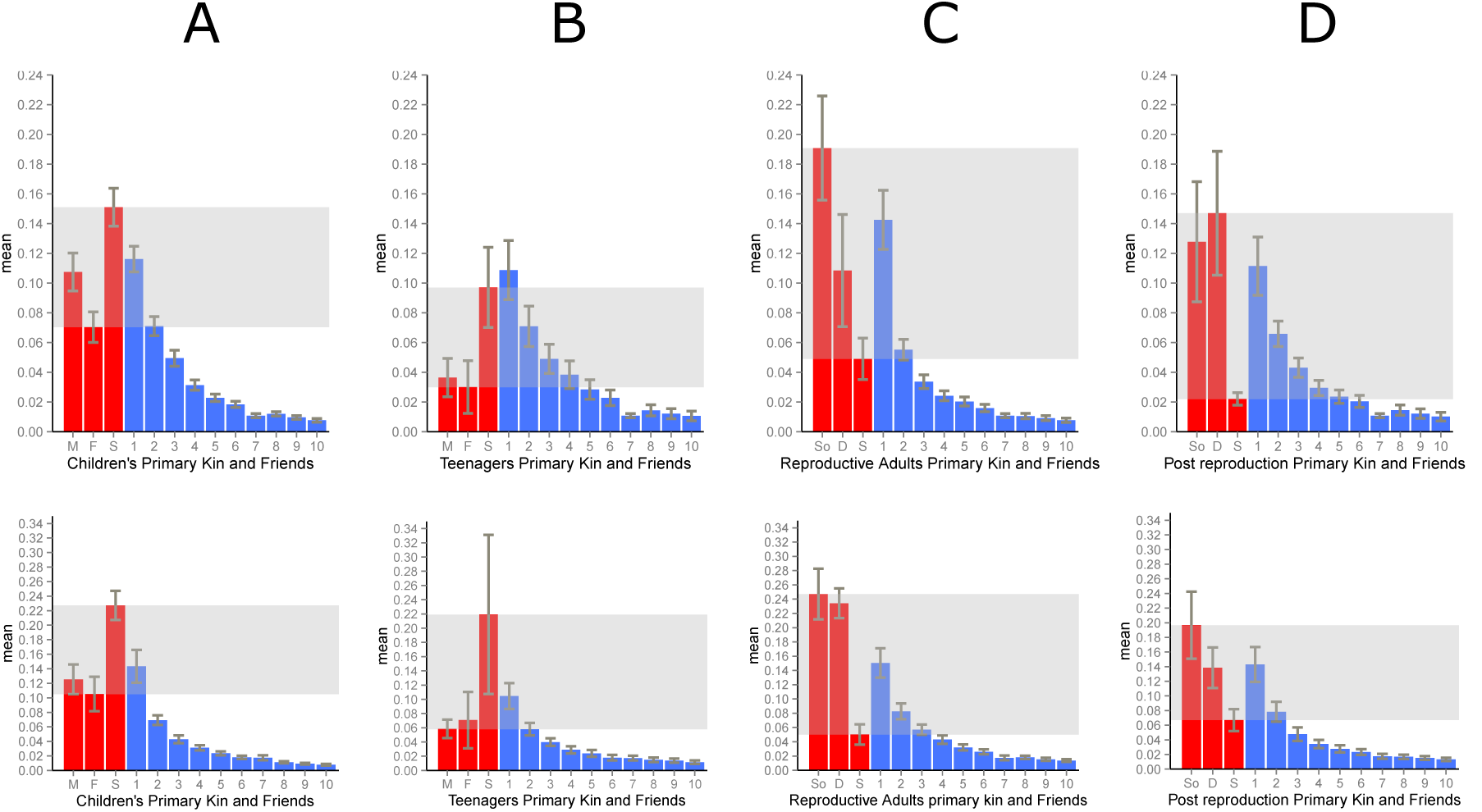
Frequency of close-range interactions with close kin and unrelated individuals. Top row, Philippines (all camps); bottom row, Congo (all camps). (A) children (2-12 years), (B) teenagers (13-17) (C) reproductive adults (18-45), (D) post-reproductive adults (46 or over). Red bars: from left to right, proportion of interactions with mother, father and siblings (A and B); or sons, daughters and siblings (C and D). Blue bars: proportion of interactions with unrelated individuals ranked from left to right by frequency of interactions, up to the 10^th^ strongest relationship. Spouses and affines were excluded. Shaded area represents the range of frequency of interactions with close kin. In all plots, error bars represent plus and minus one standard deviation. In both camps and across all age groups, people interact with from one to four unrelated individuals as closely as with their close kin.

### Human life history is adapted to the development of non-kin relationships

If non-kin interactions and investment in a few strong friends explain network global efficiency and human hypersociality, we should expect non-kin interactions to develop very early in human ontogeny as an important human life history adaptation. Children exhibit from an early age cognitive tendencies that predispose them to social norm acquisition, learning and cooperation^14, 15^, such as imitation^16^ and shared intentionality^17^. Those tendencies are extended to interactions with unrelated peers ^18^, with potential effects on the development of cooperative networks ^3^ and cultural transmission^5^. Among the Agta, 27% of interactions of children between the ages of 3 and 7 occurred with non-kin (Fig. 3), compared to 32% of interactions with siblings, 13% with mothers, and less than 1% with their grandmothers. Among BaYaka, 30% of interactions of post-weaning children aged 2 to 7 were with non-kin (Fig. 3), 30% with siblings, 17% with mothers, and 5% with grandmothers. Between ages 8 and 12, interactions with non-kin increase to 39% in the Agta and 35% in the BaYaka. Overall, non-kin interactions among children aged between 2 and 12 years show significant age-assortativity (Philippines: β=26.6, P < 0.001, 95% CI = [14.6, 38.67]; Congo: β=29.3, P < 0.001, 95% CI [18.7, 38.8]; see Methods). The high frequency of non-kin relations at early ages has many potential implications, allowing a unique involvement of unrelated individuals in allocare, creating opportunities for social learning ^19^ and learning through simulation of adult behaviours in child-only play groups (see Supplementary Video 1), in addition to laying the foundations for adult social networks ^20^.

**Figure 3.**
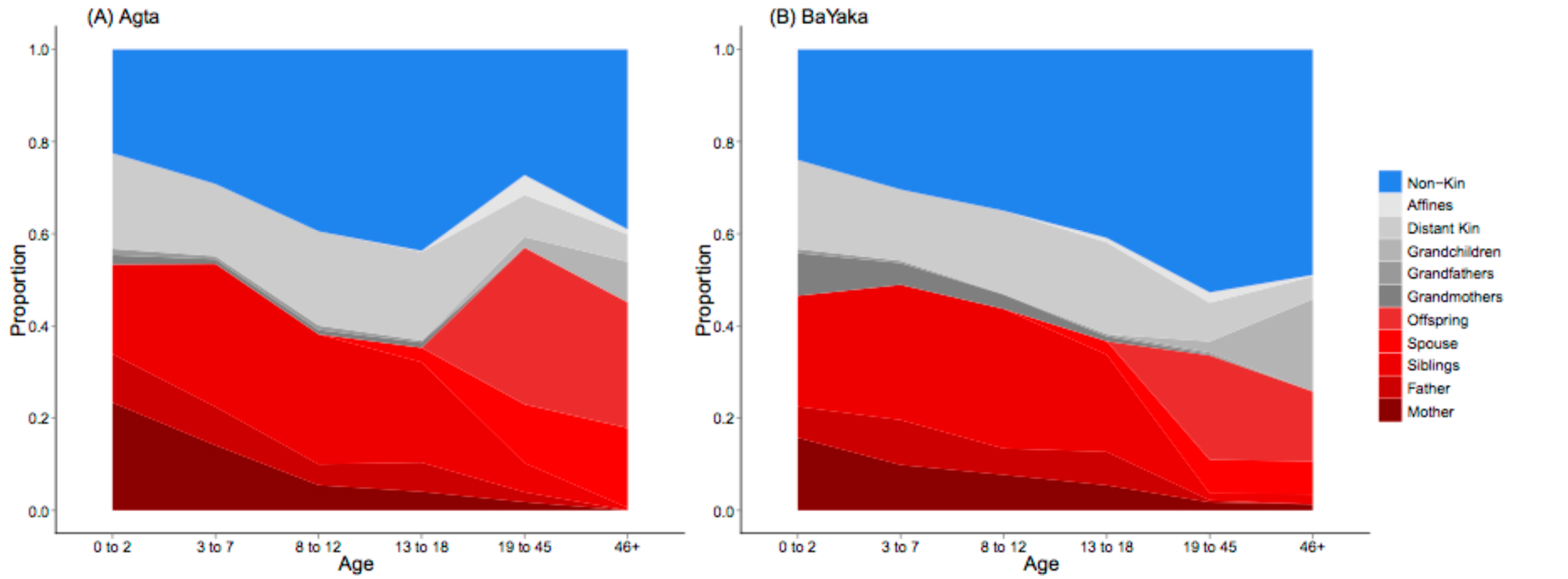
Proportion of interactions by age group and relatedness category. Colours represent relatedness categories (close kin: mother, father, siblings, spouse, offspring; extended family: grandparents, grandchildren, aunt, uncle, niece, nephew, first cousins, parents-in-law, siblings-in-law; non-kin: all other individuals). (A) Philippines, all camps. (B) Congo, all camps. From an early age, weaned children (aged 2-7) exhibit a large frequency of interactions with unrelated individuals in play groups (see main text).

## Discussion

The early developmental origin of human socio-cognitive abilities to establish links with non-kin has important implications for our understanding of human life history. Surprisingly, in both Agta and BaYaka around 30% of recorded interactions of children aged 2 and 7 were with non-kin. We argue that the human unique slow life history and delayed maturation are not only a consequence of high energetic offspring costs ^21^ and intergenerational transfers ^22^, but also an adaptation that facilitates social learning through cultural diffusion in play groups, where children are frequently looked after by older children, learn through playing and imitation of role models (see Supplementary Video 1), cooperate with same-aged children ^19^, attempt innovations and acquire sociocultural information from unrelated children ^23^.

In Agta and BaYaka play groups children also establish their first friendships, which accomplish an important role in adult life. We show that across age groups people have at any given time on average one to four ‘best friends’, and a large number of less intense links with other unrelated individuals. These friendships are likely to be one of the conditions for the high between-camp mobility that characterise hunter-gatherers ^24^, who encounter around ten times more individuals over a lifetime than chimpanzees ^25, 26^. We propose that the continuous movement of friends may create an infrastructure of networks across multiple camps, facilitating cultural flows and assortment on the basis of genetic or behavioural cues such as the tendency to cooperate ^3^.

Non-kin ties are also central to the optimisation of information flow across human networks. Given the practical impossibility of cultivating strong social bonds with all unrelated individuals, hunter-gatherers focus on a few important friendships among a large number of acquaintances. Our results show that the global efficiency of hunter-gatherer networks depends on a combination of weak^27^ and strong non-kin ties, and is drastically reduced in randomised networks which lack strong interactions with ‘best friends’. It is important to notice that efficient networks may also have disadvantageous consequences, such as the spread of infectious diseases, which might have devastating consequences for hunter-gatherers due to their often low population sizes. However, real-world networks are known to be dynamic and adapt to the state of particular nodes by breaking ties and temporarily reducing efficiency when required ^28^. In highly mobile hunter-gatherers such as the Agta, we observed an example of adaptive rewiring of proximity networks in one camp, which broke down into two smaller units that moved away from each other during an outbreak of measles.

Optimised network efficiency in human groups may have been fundamental for the evolution of human unique characteristics, such as the sharing of food, technology and ideas, group-level cooperative actions and especially the evolution of cumulative culture. In hunter-gatherers, extreme camp fluidity, multilocality and the egalitarian socio-political structure allow households to frequently move camps, resulting in co-residence with a large number of unrelated individuals ^29,30^. Establishing non-kin links may be therefore a condition for information to be shared and accumulated more widely across camps. As social groups evolved from small hunter-gatherer camps into larger groupings with the advent of agriculture, and later into our present megacities, the opportunities for people to extend their social networks and increase links with unrelated individuals also expanded. We have shown that the developmental and behavioural adaptations which make this possible had already been established in small-scale, nomadic societies. We propose that such adaptations explain why people are keen to socialise, cooperate and exchange information with unknown individuals in megacities and even in global-scale social networks on the World Wide Web.

## Methods

### 1. Sample

We studied two populations of hunter-gatherers: the Agta from the Philippines, and the Mbedjele BaYaka pygmies from Congo between March and September 2014.

#### 1.1. Agta

The Agta are hunter-gatherers from the Philippines who subsist on terrestrial, river and costal marine resources. They live in North East Luzon, within the Northern Sierra Madre Natural Park, Municipality of Palanan, Isabela and speak Agta Paranan (an Austronesian Language). The Agta population is estimated to be around 1000 individuals in the Palanan region. We studied 200 individuals of all ages from six camps. They live in small bands of 49±22 people an average. There is a range of mobility within the population as some camps are comprised of semipermanent houses and others are more fluid, with some households moving between camps more regularly than others. The Agta trade some of their fish and forest products for rice and occasionally engage in cash labour.

#### 1.2. Mbendjele BaYaka

The Mbendjele are a subgroup of the BaYaka Pygmy hunter-gatherers who speak Mbendjele (a Bantu language). Their residence spans across the forests of Congo-Brazzaville and Central African Republic, and the total population in that region is estimated in 30,000. The study population lives in the marsh rainforests of the Sangha and Likuoala regions. BaYaka subsistence techniques include hunting, trapping, fishing, gathering and honey collecting. The BaYaka live in langos—multi-family camps consisting of a number of fumas (huts) in which nuclear families reside. Camp size tends to vary from 10 to 60 individuals, with an average of 44±24 people. We studied 132 Mbendjele of all ages distributed in three camps. The study population trade some of their meat and forest products for farmer products and occasionally engage in cash labour. They live in the forest or near the mud roads opened by logging companies and move between camps opportunistically depending on food resources and trade opportunities.

### 2. Portable wireless sensing technology (motes)

#### 2.1. The motes

Recent progress in embedded electronics has led to compact (50 mm*35 mm*15 mm with casing) and affordable wearable devices with sensors. For this study, we considered many solutions before selecting devices supporting TinyOS, an operating system developed at the University of California, Berkeley. Our selected device is based on the UCMote Mini with some custom modifications. It comprises of a main processor, a wireless communication module, a memory storage unit and a battery which allows the devices to run for up to four weeks in one charge once the software is optimised for low energy consumption (see Supplementary Fig. 3). We deployed 200 motes in the Philippines and 200 in Congo.

#### 2.2. The software

We wrote the embedded software in *C* and *nesC* following an iterative process with many testing phases to adjust the parameters (frequency of beacons, strength of wireless communications, and length of sleep phases) to their optimum values. In our application, each device sends beacons at a frequency of 2 minutes, receiving beacons from all other devices within a 3-meter range and storing them in long-term memory. At the end of the experiment device’s memory was downloaded via a PC side application written in JAVA.

#### 2.3. Range and calibration

The radio links were adjusted to allow a mote to record all other radio signals within a radius of approximately 3 meters. A specific radio transmission technique (low power listening) was used to reduce battery usage whilst assuring a good delivery of all messages. We calibrated radio links by testing devices on a range of situations and environments, in the UK and in the field.

#### 2.4. Motes utilisation in the field

After waterproofed with cling film motes were sealed into wristbands or armbands (for babies). We studied one camp at a time in the Philippines and Congo. After explanation methods and discussing data anonymity through presentations and posters in their local languages, we gave one mote to each participant who agreed to take part in the experiment and signed informed consent forms. Each mote was labelled with a unique number and identified with a coloured string. All individuals within a camp (from newborns to elderly individuals) wore their motes from four to nine days depending on the camp. Although motes were worn throughout the night, only data collected between 05:00 and 20:00 were analysed. If individual arrived at camp during the experiment they were promptly given a mote, and entry time was recorded. Similarly, if individual left camp at any time before the end of the experiment the time they returned the mote was recorded. A small compensation (usually a thermal bottle or cooking utensils) was given to each participant when the mote was returned at the end of the experiment. To ensure swaps did not occur individuals were regularly (often during interviews) asked to check they were wearing the correct armband. Mote numbers were also checked when motes were returned to ensure they had not been swapped. Any alterations were recorded and adjusted in the final data processing.

#### 2.5. Ethical approval

This research and fieldwork was approved by UCL Ethics Committee (UCL Ethics code 3086/003) and carried out after informed consent was obtained from all participants.

#### 2.6. Data recovery

Raw data were run through a stringent data processing system written in *Python* to leverage the filtering power of MySQL databases. The procedure ensured that the used data were free from corruption due to device damage. Once the basic checks were passed, data were matched with individual IDs number (protecting the real names) and with start and stop times of each mote. The next step was to combine all data in a database. The result was a matrix containing the number of recorded beacons for all possible dyads (i.e. the frequency of close-range interactions between pairs of individuals) for each camp. For dyads involving individuals receiving motes after the experiment had started, a correction proportional to the duration of their participation was applied.

#### 2.6. Motes validation

To establish whether or not the motes were in fact recording proximity within a three-meter range, we compared data from motes to observational data from eight children aged between 3 and 5 years old. We conducted ‘focal follows’ of a child for a total of nine hours over three non-constitutive days, observing and recording all individuals present within three meters of the child every 30 seconds^31^. This produces 1080 observational points per child over three days (one every 30 seconds), compared to an average of 3150 emitted motes points over one week (1 every 2 minutes). However, since multiple ties are captured with each observation or every two minutes with the motes, respectively, there is on average 3850 mote points captured compared to 3080 observational points per child. Focal follows are a much more invasive and intensive form of data collection and do not provide the full range of interaction of the whole population.

To compare motes and focal follows data, we produced averages for the proportion of time eight infants spent with specific kin categories. The differences between the averages of interactions for a week (motes) and observations for a day of focal follows are minimal, and the distribution of observations with specific kin types is not significantly altered between the two methods. For instance, for the eight children with ages between 3 and 5 years old, the motes recorded an average 34% of time spent with mothers, 11% with fathers, 24% with siblings and 6%, 7% and 23% for grandparents, other kin (0.125≤ r ≤ 0.25) and non-kin (r < 0.125), respectively. These same children were observed during the focal follows to spend 37% of time within three-meters of their mothers, 19% with fathers, 24 % with siblings and 2 %, 7% and 24% of their time with grandparents, other kin and non-kin, respectively. Therefore, the two types of data collection produce remarkably similar pictures of proximity. The small differences are most likely caused by motes data representing a full week, while focal follows represent only nine hours of data. Overall, the consistency between the observational and motes data lead us to conclude motes have a high reliability and represent proximity at approximately three meters. Note that the total proportions do not add up to 100% as multiple people can be found simultaneously within the three-meter range.

### 3. Genealogical data and kin definition

We collected genealogies over three generations for all individuals in the study, and built matrices of relatedness based on kin categories (mother, father, son, daughter, spouse, brother, sister, uncle, aunt, niece, nephew, cousin, grandparents, grandchildren, parents-in-law, children-in-law, brother/sister-in law, other kin, other affines, and unrelated individuals).

For the network analyses we defined ‘primary kin’ as parents, children, siblings and partners. ‘Extended family’ included distant kin (grandparents, grandchildren, aunt, uncle, niece, nephew, first cousins, parents-in-law, siblings-in-law). ‘Unrelated individuals’ include all other individuals.

### 4. Multi-level modelling of age assortativity

We tested for age assortativity in dyadic interactions using multilevel analysis implemented as a mixed-effects model. To control for pseudoreplication we defined dyad, ego ID and camp as hierarchically structured random effects, and ‘same age’ as a binary (yes/no) fixed effect. To create this variable, each individual was allocated an age group: infant (under 2 years old); child (2-12 years); teenager (13-18 years); reproductive adults (18-45 years); and post-reproductive adults (46 and over).

### 5. Social Network Analysis

We used proximity data to build ten undirected weighted graphs G describing the social interaction networks for each of the ten studied camps. The N nodes of each network represent the individuals in the camp, while the undirected link (i,j) between node i and j indicates the presence of proximity interactions between individual i and individual j. The weight w_ij_ of link (i,j) was defined by the frequency of interaction between two individuals, measured by the number of recorded interactions (beacons) between the two corresponding motes. The weights range from the smallest possible non-zero value of w_ij_=238 to w_ij_=20,876 beacons (i.e. time units of 2 minutes each). The graphs are described by the number of nodes N, and by the N x N symmetric and weighted adjacency matrix W={w_ij_}, with i,j=1,2,…,N. Entry w_ij_ is equal to zero if individuals i and j had no close-range social contacts, and by definition also when i=j. For each graph, an unweighted adjacency matrix A={a_ij_}, with i,j=1,2,…,N, can be defined by setting a_ij_=1 if w_ij_ is different from zero, and a_ij_=0 otherwise. The total number of links in the graph is equal to 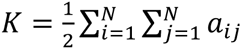. The degree k_i_ of a node i is defined as 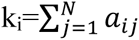, and is equal to the number of its first neighbours, while its strength s_i_ is equal to the sum of node weights 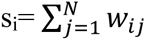. Finally, the average node degree is <ki>=2K/N.

#### 5.1. Link weight distribution and Gini coefficient

The heterogeneity in the distribution of weights among the links of a graph can be quantified by means of the Gini coefficient. The Gini coefficient g is an index used in economics and ecology to measure inequalities of a given resource among the individuals of a population^32^. It is obtained by comparing the Lorenz curve of a ranked empirical distribution, i.e. a curve that shows, for the bottom *x*% of individuals, the cumulative percentage *y*% of the total size which they have, with the line of perfect equality. In our case, we obtain the Lorenz curve by plotting the percentage *y*% of the total weights held by the *x*% of links considered, sorted in increasing value of weights. The Gini coefficient g ranges from a minimum value of zero, when all individuals are equal, to a theoretical maximum value of 1 in a population in which every individual except one has a size of zero.

#### 5.2. Calculating network efficiency

Network global efficiency of graph G was calculated as follows. First, weighted shortest paths were computed for each couple of nodes in G, assuming that the length l_ij_ of an existing link (i,j) is equal to the inverse of the weight w_ij_, and using standard algorithms to solve the all-shortest-path problem in weighted graphs. The distance d_ij_ between nodes i and j is defined as the sum of the link lengths over the shortest path connecting i and j. The efficiency ɛ_ij_ in the communication from i to j over the graph is, then, assumed to be inversely proportional to the shortest path length, i.e. ɛ_ij_=1/d_ij_. When there is no path linking i to j we have d_ij_=+∞ and, consistently, the efficiency in the communication between i and j is set equal to 0. The global efficiency of graph G is defined as the average of ɛ_ij_ over all couples of nodes:

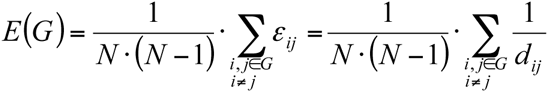

In the case of unweighted graphs, the efficiency E assumes values from 0 to 1, while in weighted graphs the values of E(G) depend on the typical weights associated to the links. It is therefore very useful to compare the efficiency found for a given weighted network to the efficiency of a properly randomised version of the network.

#### 5.3. Network randomisation

We have constructed randomisations for each of the nine undirected weighted graphs G describing their social interaction network. The main idea is to randomize each graph by maintaining some of its original properties, such as the total number of links, the sum of all the weights (corresponding to the sum of the durations of all contacts in the network), and the degree of each node, and then randomising the links and nodes at each level of relatedness. To that purpose we divided the ties into close kin, extended family, and lastly non-kin. Then, for each camp, we considered first a network with only close-kin links, and we compared it to its randomised versions. The randomisation procedure consists in the following two stages.

Stage A: changing the adjacency matrix of close-kin ties

1) Take a node i and one of its k_i_ close-kin neighbours, let us say node j.

2) Choose with uniform probability node l among all the nodes in close-kin relation with node i (excluding node j), and node m in the close-kin neighbourhood of node l.

3) If there are no links already between node i and node m, and between node j and node l while, at the same time, nodes i and m are close kin, and node j and l are close kin, swap the two links by connecting node i to node m and node j to node l.

4) If any of the conditions in point 3 are not verified, repeat the search in point 2 with another couple of nodes l and m, up to M times. If after M times the conditions have not been fulfilled, the link between node i and node j is left unaltered.

Stage B: redistributing weights to the new adjacency matrix

5) Each node i has a total number of beacons equal to its strength s_i_ (the sum of the weights of all its links). Each of these beacons is randomly reallocated with uniform probability to one of the k_i_ new neighbours.

Steps (1-5) are repeated for each node and for each of its links.

Second, we considered the network with close kin and extended family links, and then randomised only extended family links according to the procedure above. Finally, we considered the network with close kin, extended family and non-kin links, and randomised only non-kin links. For each of the three cases, we used M=100 iterations and we created an ensemble of 1000 randomised graphs. The average value of efficiency obtained for the ensemble of randomised graphs was then compared with that of the real networks at the three relatedness levels for each camp and each type of link.

## Acknowledgments

A.B.M. conceived the project, S.V. and A.P. designed the motes, A.B.M., M.D., J.T., A.P., D.S., G.D.S., N.C., S.V. collected data, G.D.S. provided video images from Congo, J.G.-G, V.L. performed social network analysis, J.G.-G., S.V., A.P., M.D., D.S., N.C., J.S., J.T., R.M., V.L., L.V and A.B.M. analysed the data, and A.B.M. and L.V. wrote the paper with the help from all other authors. The authors declare no competing financial interests. For data access please contact the correspondent author.

This project was funded by the Leverhulme Trust grant RP2011-R-045 to A.B.M. and R.M. R.M. also received funding from European Research Council Advanced Grant AdG 249347. We thank J. Lewis and R. Schlaepfer for help in the field. We thank Rodolph Schlaepfer and RKSmedia for producing the accompanying movies. We also thank our assistants in Congo and the Philippines, as well as the Agta and BaYaka communities.

## Supplementary Materials

Supplementary Figs. 1-3

Supplementary Table 1

Supplementary Movie 1

**Supplementary Figure 1.**
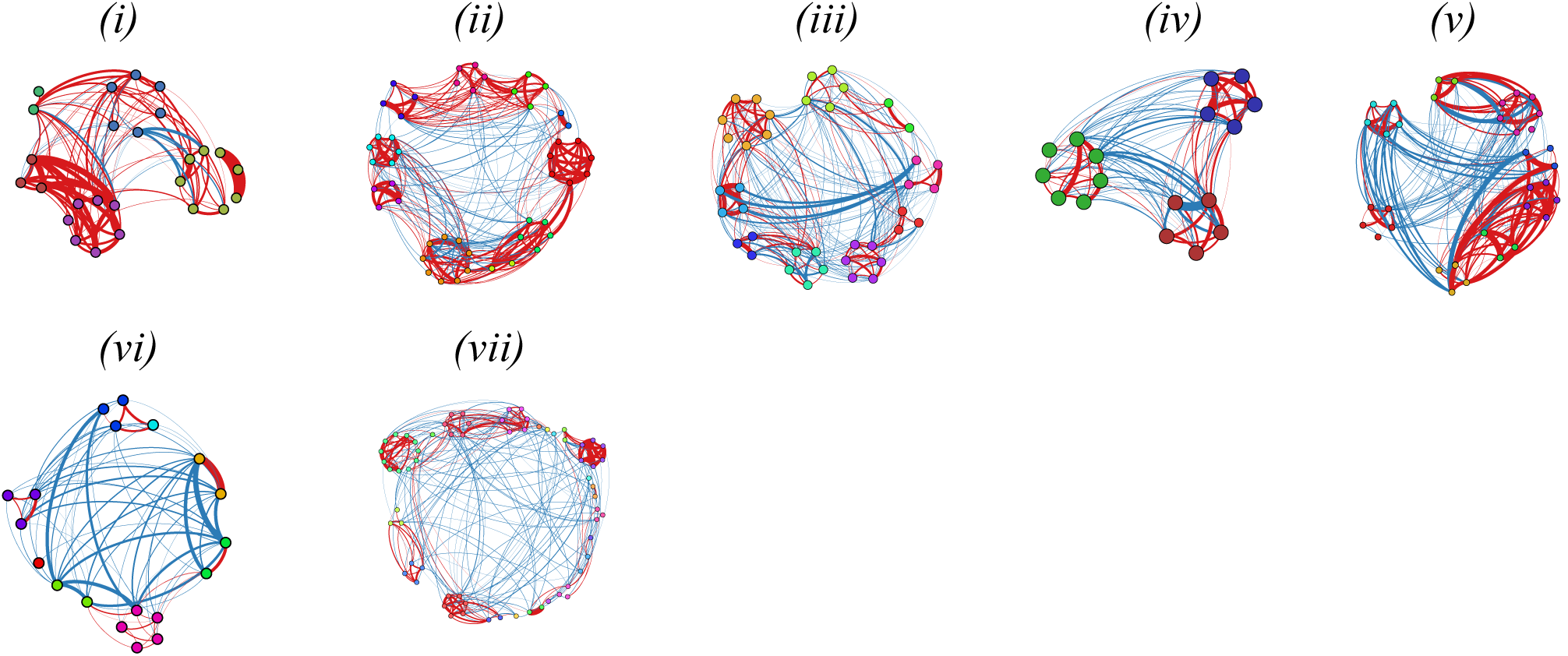
Diagrams of networks for the remaining camps in the Philippines (top 5) and Congo (bottom 2). Nodes represent individuals, node colours represent households. Red ties represent close kin and extended family, and blue ties connect unrelated individuals. Tie thickness represents intensity of relationship as measured by number of recorded close-range interactions over a week, revealing strong non-kin ties between individuals from different households. Graphs display approximately the 60% strongest links.

**Supplementary Figure 2.**
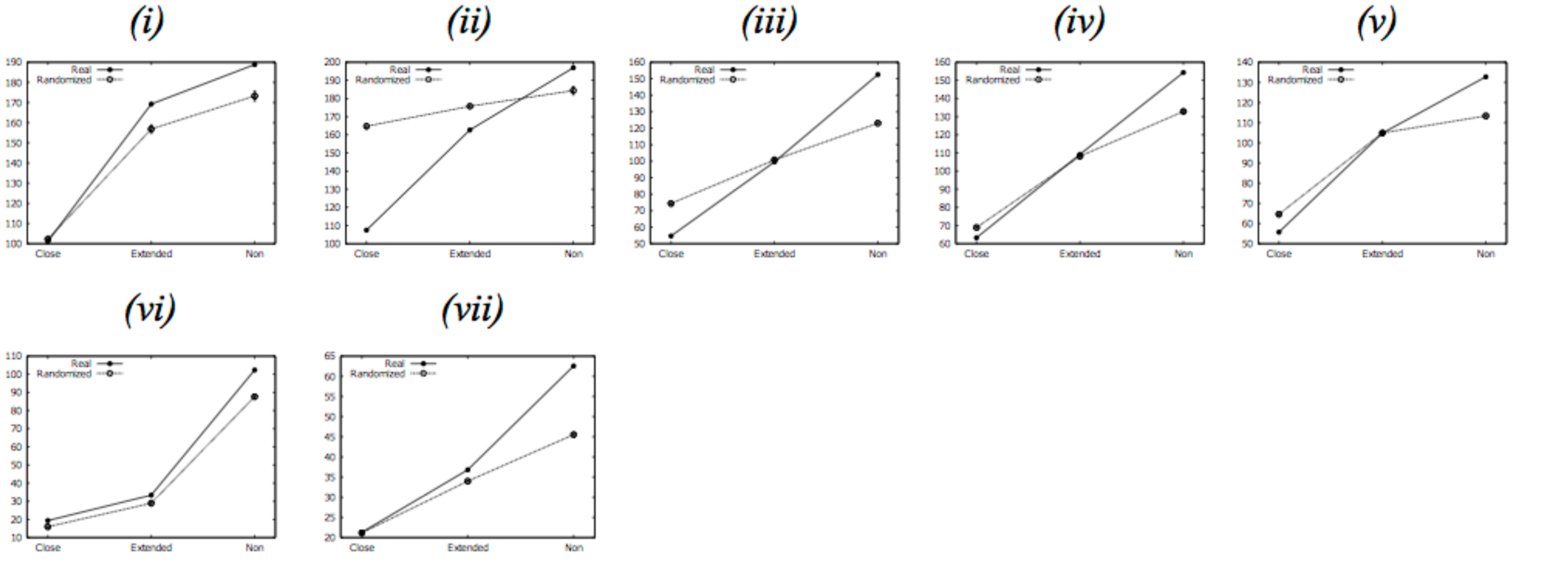
Global network efficiency in real network (solid circles) and randomised networks (open circles) for the remaining camps in the Philippines (i-v) and Congo (vi-vii). Randomisation of ties between close kin and extended family has minimal effect on global efficiency of networks. In contrast, randomisation of ties between non-kin in real networks causes dramatic reduction in global efficiency. In all comparisons, we considered averages over 1000 different randomisations. Error bars for randomisations represent standard error of mean, but are small and imperceptible.

**Supplementary Figure 3.**
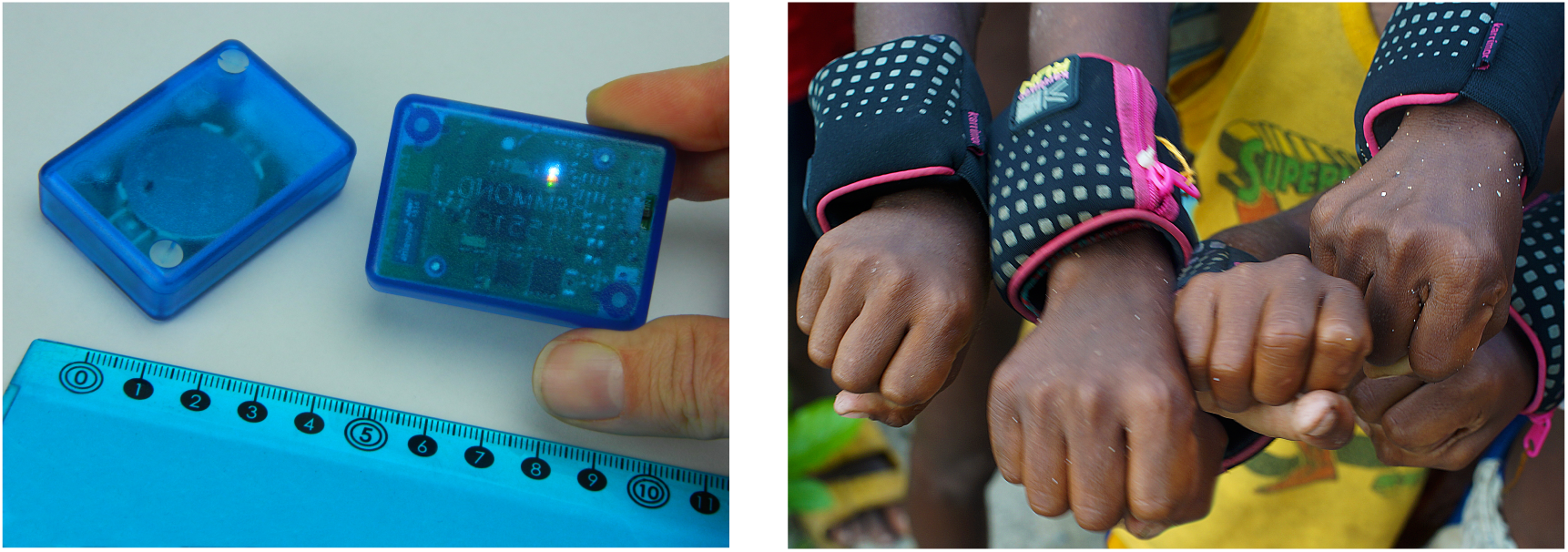
Pictures of motes (a), and of the Agta wearing them in armbands in the Philippines (b). Credit: Rodolph Schlaepfer and Sylvain Viguier

**Supplementary Video 1**. The video shows BaYaka Pygmies from Congo Brazzaville performing a forest spirits ritual called “Bobe” that lasts for hours during the night. Women sing polyphonic music and play drums with children. After some time, adult men believed to be possessed by forest spirits are attracted by the singing and come into the camp to dance. The ritual is believed to be important for group cohesion. Later the video shows children mimicking the adult performance of the forest spirits ritual in an unsupervised play group. Video footage was shot by Gul Deniz Salali between March and July 2014. Video was edited by Rodolph Schlaepfer. (RKSmedia).

## References and Notes

1. Fehr, E. & Fischbacher, U. The nature of human altruism. Nature 425, 785–791 (2003)

2. Kingma, S.A. Santema, P., Taborsky, M. & Komdeur, J. Group augmentation and the evolution of cooperation. Trends Ecol. Evol. 29, 476–484 (2014)

3. Apicella, C. L., Marlowe, F. W., Fowler, J. H., Christakis, N. A. Social networks and cooperation in hunter-gatherers. Nature 481: 497–501 (2012)

4. Burkart, J. M., Hrdy, S. B. & van Schaik, C. P. Cooperative breeding and human cognitive evolution. Evol. Anthopol. 18, 175–186 (2009)

5. Dean, L. G., Kendal, R. L., Schapiro, S. J., Thierry, B. & K. N. Laland. Identification of the Social and Cognitive Processes Underlying Human Cumulative Culture. Science, 335, 1114–1118 (2012)

6. Hamilton, M. J., Milne, B. T., Walker, R. S., Burger, O. & Brown, J. H. The complex structure of hunter–gatherer social networks. Proc. R. Soc. Lond. B 274, 2195–2203 (2007)

7. Jaeggi, A.V. & Gurven, M. Natural cooperators: food sharing in humans and other primates. Evol. Anthropol. 22, 186–195 (2015)

8. Kramer, K.L. The evolution of human parental care and recruitment of juvenile help. Trends Ecol. Evol. 26, 533–540 (2011).

9. Powell, A., Shennan, S., Thomas, M.G. Late Pleistocene Demography and the Appearance of Modern Human Behavior. Science, 324, 1298–1301 (2009)

10. Latora, V. & Marchiori, M. Economic Small-World Behavior in Weighted Networks. Eur. Phys. J. B 32, 249–263 (2003)

11. Watts, D.J. & Strogatz, S.H. Collective dynamics of 'small-world' networks. Nature 393, 440–442 (1998)

12. Wohlgemuth, J. & Matache, M.T. Small-World Properties of Facebook Group Networks. Complex Systems 23, 3 (2012)

13. Albert, R., Jeong, H., Barabási, A.-L. Diameter of the world wide web. Nature 401, 130–131 (1999)

14. Salali, G.D., Juda, M., Henrich, J. Transmission and development of costly punishment in children. Evolution and Human Behavior, 36: 86–94 (2014)

15. House, B.R. et al. Ontogeny of prosocial behavior across diverse societies. Proc. Natl. Acad. Sci. USA 110, 14586–14591 (2013)

16. Ray, E. D. & Heyes, C. M. Imitation in infancy: The wealth of the stimulus. Develop. Sci., 14, 92–105 (2011)

17. Tomasello, M. & Carpenter, M. Shared intentionality. Develop. Sci. 10, 121–125 (2007)

18. Fehr, E., Bernhard, H. & Rockenbach, B. Egalitarianism in young children. Nature, 454, 1079–1083 (2008)

19. Warneken, F., Steinwender, J., Hamann, K., & Tomasello, M. Young Children’s Planning in a Collaborative Problem-Solving Task. Cog. Dev., 31, 48–58 (2014)

20. Chaudhary, N., Salali, G. D., Thompson, J., Dyble, M., Page, A., Smith, D., Mace, R. & Migliano, A. B. Polygyny without wealth: popularity in gift games predicts polygyny in BaYaka Pygmies. R. Soc. open sci. 2, 150054 (2015)

21. Kuzawa, C. W. et al. Metabolic costs and evolutionary implications of human brain development. Proc. Natl. Acad. Sci. USA 111, 13010–13015 (2014)

22. Kaplan H & Robson. The emergence of humans: The coevolution of intelligence and longevity with intergenerational transfers. Proc Natl Acad Sci USA 99, 10221–10226 (2002)

23. Whiten A. & Flynn E. The transmission and evolution of experimental microcultures in groups of young children. Develop. Psych. 46, 1694–709 (2010)

24. Lewis, H.M., Vinicius, L., Strods, J., Mace, R. & Migliano, A. B. High mobility explains demand sharing and enforced cooperation in egalitarian hunter-gatherers. Nature Comm. 5, 5789 (2014)

25. Hill, K. R., Wood, B. M., Baggio, J., Hurtado, A. M. & Boyd, R. T. Hunter-Gatherer Inter-Band Interaction Rates: Implications for Cumulative Culture. PLoS ONE 9, e102806 (2014)

26. Dunbar, D. How Many Friends Does One Person Need? Dunbar’s Number and Other Evolutionary Quirks. Harvard University Press, Cambridge MA (2010)

27. Granovetters, M. The Strength of Weak Ties. Am. J. Sociol. 78, 1360–80 (1973)

28. Gross, T., D’Lima, C. J. D. & Blasius, B. Epidemic dynamics on an adaptive network. Phys. Rev. Lett. 96, 208701 (2006).

29. Hill, K. R. et al. Co-residence patterns in hunter-gatherer societies show unique human social structure. Science 331, 1286–1289 (2011)

30. Dyble, M. et al. Sex equality can explain the unique social structure of hunter-gatherer bands. Science 348, 796–798 (2015)

31. Meehan, C. L., Quinlan, R. & Malcom, C. D. Cooperative breeding and maternal energy expenditure among Aka foragers. Am. J. Hum. Biol. 25, 42–57 (2013).

32. Dagum, C. The generation and distribution of income, the Lorentz curve and the gini ratio Écon. Appl. 33, 327–367 (1980).

